# An evolutionary-conserved Wnt3/β-catenin/Sp5 feedback loop restricts head organizer activity in *Hydra*

**DOI:** 10.1101/265785

**Authors:** Matthias C. Vogg, Leonardo Beccari, Laura Iglesias Ollé, Christine Rampon, Sophie Vriz, Chrystelle Perruchoud, Yvan Wenger, Brigitte Galliot

## Abstract

The *Hydra* polyp regenerates its head by transforming the gastric tissue below the wound into a head organizer made of two antagonistic cross-reacting components. The activator, previously characterized as *Wnt3*, drives apical differentiation by acting locally and auto-catalytically. The uncharacterized inhibitor, produced under the control of the activator, prevents ectopic head formation. By crossing RNA-seq data obtained in a *β-catenin*(RNAi) screen performed in planarians and a quantitative analysis of positional and temporal gene expression in *Hydra*, we identified Sp5 as a transcription factor that fulfills the head inhibitor properties: a Wnt/β-catenin inducible expression, a graded apical-to-basal expression, a sustained up-regulation during head regeneration, a multi-headed phenotype when knocked-down, a repressing activity on Wnt3 expression. In mammalian cells, *Hydra* and zebrafish Sp5 repress *Wnt3* promoter activity while *Hydra Sp5* also auto-activates its expression, possibly via β-catenin and/or Tcf/Lef1 interaction. This work identifies Sp5 as a novel potent feedback loop inhibitor of Wnt/β-catenin signaling across eumetazoans.

## INTRODUCTION

The freshwater polyp *Hydra,* which belongs to Cnidaria, a sister group to Bilateria (**Figure 1A**), has the remarkable talent to regenerate any lost body parts, including a fully functional head. *Hydra,* which is made of two cell layers, external named epidermis and internal named gastrodermis, shows a polarized tubular anatomy with a head at the apical pole and a foot at the basal one, both extremities being enriched in nerve cells (**Figure 1B**). Head regeneration relies on the rapid transformation of a piece of a somatic adult tissue, the amputated gastric tube, into a tissue with developmental properties named head organizer, which directs the patterning of the regenerating tissue (reviewed in (Bode, 2012; Shimizu, 2012; Vogg et al., 2016)). This process is highly robust in *Hydra,* occurring after bisection at any level along the body column. The concept of organizer was first discovered through the pioneering work of Ethel Browne who performed lateral transplantation experiments between pigmented and depigmented *Hydra* (Browne, 1909). By grafting a non-pigmented piece of head onto the body column of a pigmented host, she observed the development of an ectopic axis predominantly made of pigmented cells, i.e. recruited by the graft from the host. This discovery was later confirmed in hydrozoans (Mutz, 1930; Yao, 1945; Webster, 1966; MacWilliams, 1983b; Broun and Bode, 2002) but also in vertebrates where organizers play an essential role during embryonic development (Spemann and Mangold, 1924). In *Hydra* regenerating its head, the organizer gets established within 10 to 12 hours after mid-gastric bisection, restricted to the head-regenerating tip within the first 24 hours, remaining stable until the new head is formed and subsequently persisting as a homeostatic head organizer (MacWilliams, 1983b).

The *Hydra* model also helped understand the dual structure of organizers. By comparing the efficiency of apical grafts to induce ectopic axis either on intact or on decapitated hosts, Rand et al. showed that the *Hydra* head organizer exerts two opposite activities, one positive named head activator, which promotes apical differentiation, and another negative named head inhibitor, maximal in the apical region, which prevents the formation of supernumerary or ectopic heads (Rand et al., 1926). In *Hydra* the inhibitory activity is graded along the body axis, with a maximum at the apical pole (Webster, 1966). It is also tightly modulated during head regeneration, when it rapidly decays after amputation, and slowly recovers thereafter (MacWilliams, 1983a). Interestingly, head inhibition was found significantly higher in head-regeneration deficient (Kemmner and Schaller, 1981; Achermann and Sugiyama, 1985) or low budding (Takano and Sugiyama, 1983) strains. Gierer and Meinhardt used the results obtained from a series of transplantation experiments to test their general mathematical model of morphogenesis (Gierer and Meinhardt, 1972). Their model is based on the reaction-diffusion model proposed by Turing where two substances that exhibit distinct diffusion properties and interact with each other, form a minimal regulatory loop that suffices for *de novo* pattern formation (Turing, 1952). Gierer and Meinhardt refined this model by posing that the activation component acts over short-range distance, while the inhibition acts over long-range distance, and by distinguishing between *"effective concentrations of activator and inhibitor, on one hand, and the density of their sources on the other"* (Gierer and Meinhardt, 1972). These models proved to efficiently simulate basic properties of pattern formation and were validated by molecular data in a variety of developmental contexts (Kondo and Miura, 2010).

In *Hydra,* the Holstein lab identified *Wnt3* as a growth factor fulfilling the criteria of the head activator, expressed locally at the tip of the head in intact *Hydra*, rapidly re-expressed in head-regenerating tips after amputation, and able to trigger an autocatalytic feedback loop (Hobmayer et al., 2000; Lengfeld et al., 2009; Nakamura et al., 2011). Concerning the head inhibitor necessary to maintain a single head in homeostatic polyps and to develop a single head in budding and regenerating *Hydra*, attempts were made to characterize it, either biochemically or genetically. A protease-resistant small hydrophilic molecule was identified, exhibiting an apical to basal graded activity although with some activity also detected in the basal disc (Berking, 1977; Schaller et al., 1979), and the characterization of this inhibitor was not pursued. A genetic screen identified a *Hydra* ortholog of the vertebrate Wnt inhibitors *Dkk1, Dkk2 and Dkk4,* named *hyDkk1/2/4,* which efficiently antagonizes Wnt activity in the zebrafish (Guder et al., 2006). Wntβ-catenin signaling negatively regulates *Dkk1/2/4* expression, therefore not expressed in the head, and a multi-headed phenotype was observed neither for *Dkk1/2/4* (Augustin et al., 2006; Guder et al., 2006) Therefore, the molecular nature of the negative regulator(s) of the *Hydra* head organizer remains unknown.

**Figure 1.**
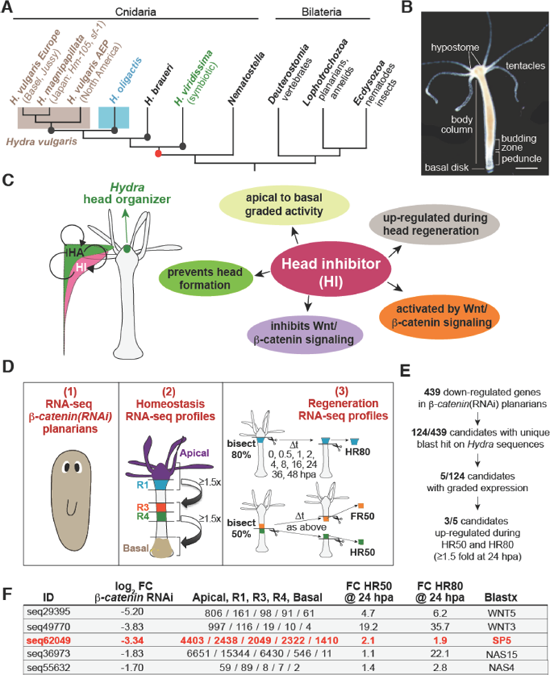
Screening strategy to identify candidate head inhibitor (HI) genes. **(A)** Tree showing the sister position of Cnidaria to Bilateria, the four *Hydra* species (black dots) and their last common ancestor (red dot). **(B)** Anatomy of an intact *Hydra.* The apical extremity (head), is composed of a dome-shaped structure called hypostome, surrounded by a ring of tentacles. At the other extremity (foot), the basal disk allows the animals to attach. **(C)** The five criteria used to identify HI candidate genes. **(D, E)** Screening procedure applied to identify HI candidate genes: A first RNA-seq dataset of 439 genes down-regulated in *β*-catenin(RNAi) planarians (Reuter et al. 2015) was used to retrieve through blastx on NCBI (E-value <1e^−10^) 124 non-redundant *Hydra* sequences that correspond to 106 unique proteins (**Table Supplement 1**). These 124 candidates were next tested on RNA-seq datasets obtained in intact *Hydra* measured at five positions along the body axis (apical -Ap-, regions R1, R3, R4, basal -Ba-) and five apical-to-basal graded genes were identified. These five candidates were then tested on RNA-seq datasets obtained from regenerating tips taken at nine time points after a 50% or 80% bisection. **(F)** Three genes down-regulated after *β-catenin* RNAi in planarians, show an apical-to-basal graded expression in *Hydra,* and a minimal 1.5 fold up-regulation in head-regenerating tips at 24 hpa. FC: Fold Change.

In this study, we initially established a list of five criteria that *Hydra* head inhibitor candidates would have to fulfill (**Figure 1C**). Next we designed a screening strategy where we crossed quantitative transcriptomic data obtained in two evolutionarily-distant species, both with high regenerative potential, the planarian worm *Schmidtea mediterranea* and the freshwater hydrozoan polyp *Hydra vulgaris,* to select candidate inhibitors of apical patterning. By testing in *Hydra* the spatial and temporal gene regulation of a dataset of putative Wnt/β-catenin target genes obtained in planarians (Reuter et al., 2015), we characterized the zinc finger transcription factor Sp5 as the most promising head inhibitor. Indeed we show here that Sp5 acts as a transcriptional repressor of *Wnt3,* leading to a robust multiheaded phenotype when knocked-down in *Hydra,* while *Hydra Sp5 (HySp5)* expression is positively modulated by Wnt/β-catenin signaling in *Hydra* as well as in mammalian cells. Thus Sp5 fulfills the requirements of a head inhibitor in *Hydra*, and we also show that the feedback loop function of Sp5 on Wntβ-catenin signaling appears conserved across eumetazoan evolution.

## RESULTS

### Identification of putative head inhibitors by crossing RNA-seq datasets from planarians and *Hydra*

To identify head inhibitors, we reasoned that putative candidates should (1) be activated by Wnt/β-catenin signaling, (2) exhibit an apical to basal graded activity in intact animals, (3) be upregulated within the first 24 hours of head regeneration, (4) inhibit Wnt/β-catenin signaling, (5) prevent head formation (**Figure 1C**). To select candidates activated by Wnt/β-catenin signaling, we used a dataset of 439 genes that are down-regulated in planarians silenced for *β-catenin* (Reuter et al., 2015) (**Figure 1D**, left). Next we identified among these putative Wnt/β-catenin target genes 124 besthit blastx genes expressed in *Hydra* (**Table Supplement 1**). When tested for their spatial homeostatic expression through RNA-seq profiles measured at five distinct positions along the *Hydra* body axis (**Figure 1D**, center), we found 5/124 candidates with an apical to basal graded expression pattern, possibly reflecting a graded activity (**Figure 1E, 1F – Table Supplement 1**). The analysis of the expression of these candidate genes during regeneration, measured by RNA-seq at nine different time-points in three distinct regenerative contexts (HR80, HR50, FR50 as defined in **Figure 1D**), showed that 3/5 are up-regulated at least 1.5 fold at 24 hours post-amputation (hpa) (**Figure 1E, F**). These three candidates encode Wnt3, that fulfills the criteria of head activator (see above), Wnt5, that positively contributes to evagination processes (Philipp et al., 2009) and the transcription factor Sp5, expressed at highest levels in the apical region, previously identified as a Wntβ-catenin target gene in developing vertebrates (Thorpe et al., 2005; Weidinger et al., 2005; Fujimura et al., 2007; Park et al., 2013; Kennedy et al., 2016) (**Figure 1F – Figure Supplement 1**). We thus decided to characterize the putative role of Sp5 on *Hydra* head organizer activity.

### *Hydra* Sp5 (HySp5), an evolutionarily-conserved transcription factor

Sp5 is a member of the Sp/Klf class of transcription factors that bind DNA through their three Cys2/His2-type zinc fingers (ZF) located at the C-terminus (**Figure Supplement 2A**). Sp transcription factors also contain a buttonhead (BTD) box located upstream to the DNA-binding domain and involved in transactivation, and at the N-terminus a short SP box with an unknown function (Zhao and Meng, 2005). These three domains are well conserved in HySp5 (**Figure Supplement 2B**), which in phylogenetic analyses groups with the bilaterian Sp5 sequences (**Figure Supplement 3**). In fact the two major Sp families identified in bilaterians, Sp5 and Sp6-9 (Schaeper et al., 2010) can be traced in cnidarians, whereas the unique Sp sequence identified in *Amphimedon queenslandica* cannot be affiliated to any of these, and no typical Sp sequence could be found in non-metazoan species. Therefore a unique *Sp* gene arose at the base of metazoans to diversify in eumetazoan ancestors.

### *Sp5* is predominantly expressed at the apex in *Hydra*

Next we validated the *Sp5* RNA-seq profiles (**Figure 2A**) by whole mount *in situ* hybridization (WISH), confirming the apical to basal graded expression in intact *Hydra* with a maximal *HySp5* expression in the apical region although lower at the tip where *Wnt3* expression is maximal (**Figure 2B, 2C** – **Figure Supplement 4A, 4C**). This graded pattern, well visible in growing buds, is not ubiquitous as in numerous animals *Sp5* expression is strong along the central body column, bordered by two negative stripes on each side, i.e. in the upper body column and in the peduncle region (see animals 1-4 in **Figure Supplement 5)**. We also analyzed by RNA-seq the cell-type expression of *Sp5* and *Wnt3,* and found both genes predominantly expressed in the gastrodermal epithelial stem cells (gESCs), a cell type associated with morphogenetic processes (see Vogg et al. 2016) (**Figure Supplement 6**). *Sp5* transcripts are also detected in the epidermal ESCs -eESCs-although less abundant, and even less in the interstitial stem cells (ISCs). By contrast, *Wnt3* expression is restricted to gESCs.

### Up-regulation of *Sp5* in head-regenerating tips

After mid-gastric bisection, *Sp5* is up-regulated in both head- and foot-regenerating tips before 8 hpa. However its expression vanishes at 12 hpa in foot-regenerating tips but remains sustained in the head-regenerating tips (**Figure 2B** – **Figure Supplement 4B-C**). The graded *Sp5* expression pattern is well visible in head-regenerating halves, while foot-regenerating halves transiently show two poles of high *Sp5* expression, one in the head region, seemingly not affected by the bisection, and the second, intense in the foot-regenerating tips. The regions adjacent to these two poles do not express *Sp5*, generating a transient “stripe pattern” observed at 8 and 12 hpa, as a five stripe pattern in large animals. At 24 hpa, *HySp5* expression is no longer detected in most foot-regenerating tips but sustained in head-regenerating ones. Such distinct *Sp5* regulations in head- and foot-regenerating halves support the idea that *Sp5* is involved in head but not foot regeneration.

### *Hydra* Sp5, a robust head inhibitory component

Next we silenced *HySp5* by electroporating siRNAs in intact animals. Within two days following the third electroporation (EP3), HySp5(RNAi) animals started to develop ectopic axes, initially from the budding zone, few days later from the upper body column (**Figure 2C – Figure Supplement 7A**). These axes differentiate multiple heads that all express the apical markers *Wnt3* and *HyBra1* as well as the neuropeptide RF-amide, revealing their highly organized nervous system (**Figure 2C**). This multiheaded phenotype is robust and emerges quite synchronously, recorded in 50% animals 24 hours post-EP2 and in 100% 48 hours post-EP3 (**Figure 2D – Figure Supplement 7B**). To test the functionality of these ectopic heads, we fed HySp5(RNAi) animals with live *Artemia* and observed a normal feeding behavior, i.e. these heads catch and ingest the swimming preys (**Figure 2E** – **Movie 1, 2**).

To test whether HySp5(RNAi) animals regenerate multiple heads, we knocked-down *HySp5* by two successive electroporations and bisected at mid-gastric position 24 hours after EP2 when animals are still single-headed. *HySp5{*RNAi) animals regenerate multiple heads **(Figure 2F),** which contain clustered apical cells expressing *Wnt3* and exhibit a functional feeding behavior **(Figure 2F, 2G – Movie 3, 4).** Altogether, these results demonstrate that HySp5 acts as a strong head inhibitory component.

**Figure 2.**
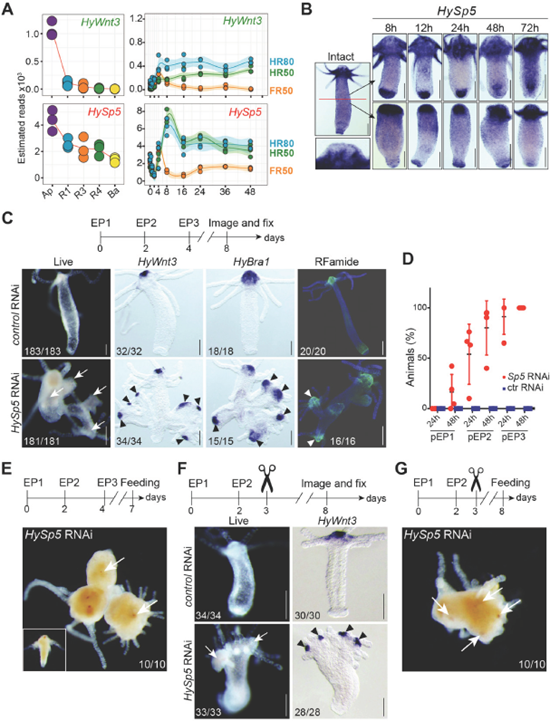
HySp5 behaves as a head inhibitory component. (A) *Wnt3* and *HySp5* RNA-seq profiles in homeostatic and regenerating animals. (B) *HySp5* expression pattern in intact and regenerating *Hydra* at indicated times after midgastric bisection (n>20, 2 experiments). Note the low *HySp5* expression in the most apical area of heads (inset below intact animal) and the sustained versus transient expression in head-and foot-regenerating tips respectively. (C) Multiheaded phenotype (white arrows) observed in intact HySp5(RNAi) *Hydra* either pictured live or fixed on day four after the third electroporation (EP3) and tested for *HyWnt3, HyBra1* (black arrowheads) or RFamide (white arrowheads) expression. (D) Kinetics of observed phenotypic changes. Each dot represents an independent experiment (n= 4). (E) Feeding response tested in HySp5(RNAi) intact animals on day-3 after EP3. White arrows point to *Artemia* ingested by the ectopic heads *(Artemia* shown in inset, orange dots correspond to the eyes). (F) Multiheaded phenotype (white arrows) observed in head-regenerating HySp5(RNAi) *Hydra* bisected at mid-gastric level 24 hours after EP2 (midgastric bisection), pictured five days later and detected for *HyWnt3* expression (black arrowheads, two experiments). (G) Feeding response tested in heads regenerated from HySp5(RNAi) animals. White arrows point to *Artemia* ingested by the multiple heads. Scale bars: 250 μm (B): 200 μm (C.F).

**Figure 3.**
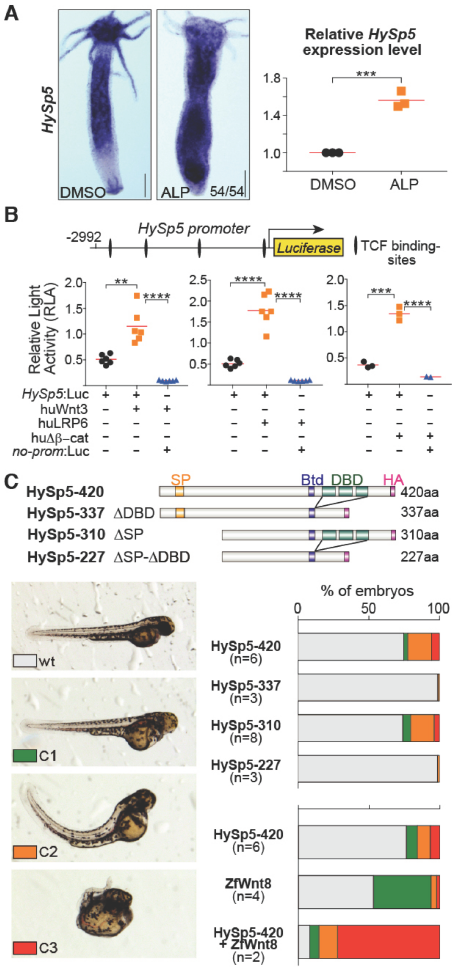
Figure 3. HySp5 responds to Wnt/β-catenin signaling in *Hydra* as in mammalian cells, and phenocopies Wnt-like phenotypes in zebrafish embryos. (A) *HySp5* expression in intact *Hydra* treated with Alsterpaullone (ALP) for two days, detected by WISH (left) and by qPCR (right). Each point represents an independent replicate. Scale bar: 250 μm.**(B)** Luciferase activity driven by the *HySp5* promoter *(HySp5:* Luc) measured in human HEK293T cells when positive regulators of Wnt/β-catenin signaling are coexpressed: human Wnt3 (huWnt3), human LRP6 (huLRP6) or constitutively active human β-catenin (huΔβ-cat). Each data point represents one biological replicate. Statistical p-values were deduced from a student’s t-test: ∗∗ ≤ 0.01; ∗∗∗ ≤ 0.001; ∗∗∗∗ ≤ 0.0001. (C) Wnt-like phenotypes detected In 2 days old zebrafish larvae overexpressing HySp5 (HySp5-420). These phenotypes were scored in three classes: no eyes (C1); no eyes + curly axis (C2); no eyes, underdeveloped axis and curly tail (C3). The HySp5 constructs lacking the DNA-binding domain do not affect embryonic development,whereas co-injecting ZfWnt8 with HySp5-420 increases the phenotypic penetrance. The number of independent experiments (n) is indicated for each construct and the graphs show one representative experiment.

### Wnt/β-catenin signaling regulates *HySp5* expression

To test whether *Sp5* is a target gene of the Wnt pathway in *Hydra,* we pharmacologically inhibited GSK-3β, a negative regulator of Wnt/β-catenin pathway by treating the animals with the kinase inhibitor Alsterpaullone (ALP) (Leost et al., 2000) that constitutively activates Wnt/β-catenin signaling (Broun et al., 2005; Rentzsch et al., 2005). We found *HySp5* up-regulated within two days in the body column of ALP-treated *Hydra* that form ectopic tentacles from day-4 **(Figure 3A** – **Figure Supplement 5),** suggesting that *HySp5* expression reflects the level of Wnt/β-catenin signaling. In addition, we cloned the *HySp5* promoter that harbors four consensus TCF binding sites (5’-(A/T)(A/T)CAAG-3’) into a luciferase reporter construct (HySp5:Luc) and tested its activity in the human HEK293T cells. When *HySp5*:Luc was co-transfected with constructs expressing either the human Wnt3, or the human Wnt co-receptor LRP6 or a constitutively active form of human β-catenin *(CMV.*huAp-cat) (Melotti et al., 2014), we recorded a significantly enhanced luciferase activity **(Figure 3B),** indicating that Wnt/β-catenin signaling up-regulates *HySp5* expression through its promoter. Altogether these results suggest that *HySp5* is a target gene of Wnt/β-catenin signaling.

**Figure 4.**
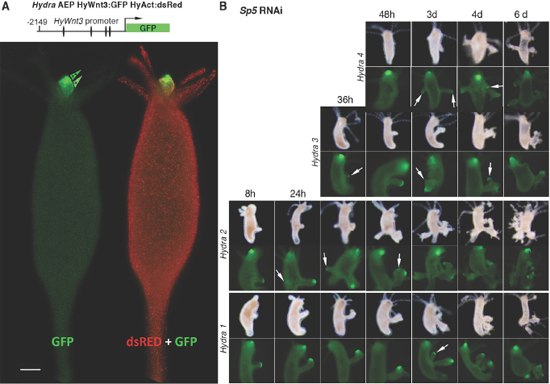
Figure 4. *Wnt3* promoter activity in homeostatic and *Sp5(RNAi) Hydra*. **(A)** Live imaging of transgenic *Hydra (Hy_AEP)* expressing GFP under the control of the *Wnt3* promoter (map shown, bars indicate TCF binding sites) and dsRED under the control of the ubiquitous *Hydra* Actin promoter (Nakamura et al., 2011). Note the two distinct levels of GFP expression in the hypostome, maximal at the tip (++), and lower (+) above the tentacle ring. (B) Live imaging of morphology (bright field) and GFP (green) expression of four *HyWnt3:GFP-HyAct:dsRED* animals knocked-down for *HySp5* with three successive siRNA electroporations. Note the GFP expression in clustered cells located at the tip of the ectopic body axes that emerge and develop along the body column (arrows).

### Overexpressing HySp5 in zebrafish embryos induces Wnt-like phenotypes

To test whether HySp5 is a mediator of the Wnt pathway, we injected *HySp5* mRNAs, either full-length (HySp5-FL) or lacking the SP box (HySp5-ΔSP) into zebrafish embryos and looked at larval morphology on day-2 post-fertilization (dpf). Overexpression of HySp5-FLand HySp5-ΔSP induced severe morphological defects ranging from no eyes (class 1 – C1), no eyes plus curly axis (class 2 – C2), to no eyes, underdeveloped axis and curly tail (class 3 – C3) (**Figure 3C**). These phenotypes are consistent with those obtained by Weidinger et al., 2005 who overexpressed the zebrafish transcripton factor Sp5a in embryos. We also found these phenotypes strongly enhanced when the zebrafish *Wnt8* mRNA was co-injected (**Figure 3C -Table Supplement 2**), but no longer present when the HySp5 DNA-binding domain was deleted (**Figure 3C**). Given the similarities with the morphological defects obtained when the zebrafish β-catenin or Wnt8 are overexpressed during development (Pelegri and Maischein, 1998; Lekven et al., 2001), we deduced that HySp5 can mediate at least some effects of Wntβ-catenin signaling during zebrafish gastrulation, a mediation that requires its DNA-binding activity.

### *HySp5* represses the activity of the *Wnt3* promoter in *Hydra*

Since *HySp5* knock-down triggers ectopic head formation in *Hydra* (**Figures 2C-G**), we postulated that HySp5 prevents head formation in *Hydra* by repressing *Wnt3.* To test this hypothesis, we produced a transgenic strain where GFP expression is directed by the *Wnt3* promoter and dsRed by the *Actin* promoter as previously tested by (Nakamura et al. 2011). These 2149-HyWnt3:GFP-HyAct:dsRed transgenic animals exhibit three levels of *Wnt3* activation in homeostatic conditions; maximal in the most apical region, intermediate in the lower hypostome region above the tentacle ring, and null at the level of the tentacle ring and along the body column (**Figure 4A**). To test *Wnt3* activation in transgenic animals knocked-down for *HySp5,* we monitored over several days after EP3 both the appearance of phenotypic traits and the GFP fluorescence (**Figure 4B**). We did not record any body-wide GFP fluorescence, which would reflect a general derepression of the *Wnt3* promoter. Instead, we noticed the successive appearance of patches of clustered GFP positive cells that take the most distal position on the ectopic axes that form along the body column. This clustered pattern of *Wnt3* derepression indicates that *HySp5* silencing is transiently efficient in clustered cells where *Wnt3* expression becomes high but without any spread GFP fluorescence in the adjacent cells. However GFP fluorescence is one day delayed compared to *GFP* expression, thus becoming visible only when ectopic axis had already developed.

### *Hy*Sp5 binds the −1/−386 *Wnt3* promoter region that represses *Wnt3*

We next assessed whether HySp5 directly regulates the *Wnt3* promoter by expressing a luciferase reporter construct driven by the *Hydra Wnt3* promoter in HEK293T cells. This HyWnt3-2149:Luc construct shows a very low basal activity, thus inappropriate to assess a putative repressive effect of HySp5. To strengthen its activity, we co-expressed a constitutively active form of β-catenin (CMV:huΔβ-cat) that enhances by ~10 fold the HyWnt3-2149:Luc activity (**Figure 5A**). When HySp5-FL is co-expressed, this β-catenin-dependent activation is significantly reduced, an inhibition not observed when the HySp5 DNA-binding domain is deleted. These results indicate that HySp5 can repress *Wnt3* promoter activity, an effect mediated through DNA-binding.

Two adjacent cis-regulatory regions were previously identified in the *Hydra Wnt3* promoter, a 599 bp long activator element that contains three clustered TCF binding sites and immediately downstream a 386 bp long repressor element (Nakamura et al., 2011). To test whether this repressor element is necessary for HySp5 repression, we produced the *HyWnt3-1763-∆Rep:Luc* construct (**Figure 5A**, right panel) and tested its activity in HEK293T cells. We found the basal activity of the *HyWnt3-1763-∆Rep:Luc* about 300× higher than that of HyWnt3-2149:Luc, still enhanced 1.5 fold when β-catenin is co-expressed (**Figure 5A**). As anticipated, HySp5 is not able to exert any repressive effect over *HyWnt3-1763-∆Rep:Luc* activity (**Figure 5A**). Noteworthy, this 386 bp repressor module and the downstream moiety of the activator element are highly conserved across *Hydra,* as shown between three evolutionarily-distant species, *H. vulgaris (Hm-105), H. oligactis and H. viridissima* (**Figure 5B**).

**Figure 5.**
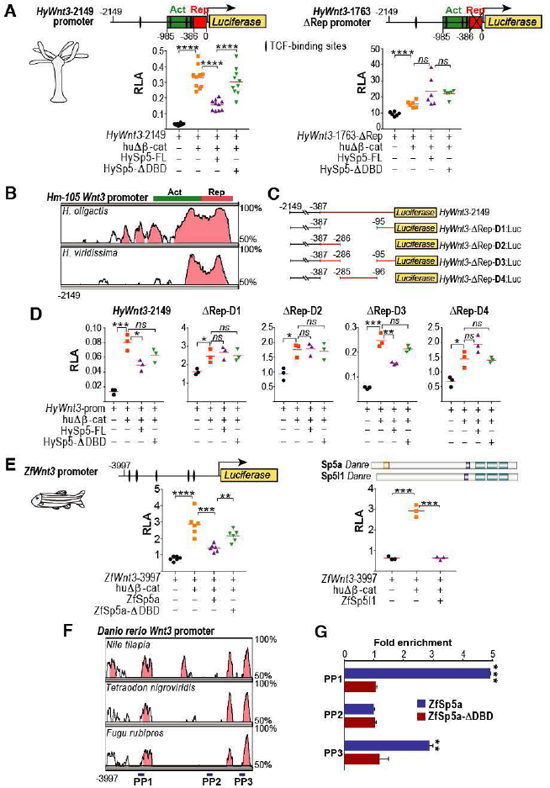
The *Hydra* and zebrafish Sp5 transcription factors repress the *Wnt3* promoter. (A) Luciferase assays measuring the activity of the HyWnt3-2149 (left) or *HyWnt3-1763-∆Rep* (right) promoters in HEK293T cells co-expressing hu∆β-cat, HySp5-FLor HySp5-ΔDBD. The Activator (Act) and Repressor (Rep) regions were identified by (Nakamura et al. 2011). Note the ~300x higher basal activity of *HyWnt3-1763-∆Rep:Luc* when compared to that of HyWnt3-2149:Luc. (B) Phylogenetic footprinting plot comparing the 2kb genomic region encompassing the *H. magnipapillata (Hm-105) Wnt3* promoter with the corresponding regions in the *H. oligactis* and *H. viridissima* genomes. Horizontal green and red bars indicate the activator (Act) and repressive (Rep) regions of the *Wnt3* promoter. Pink peaks represent evolutionary conserved modules (at least 70% base-pair identity over 100 bp sliding window). (C, D) Deletion constructs (C) used in luciferase assays performed in HEK293T cells shown in (D). Note that the repressive effect of HySp5-FL is only observed with the *HyWnt3-2149* and *HyWnt3-∆Rep-D3* constructs. (E) Luciferase assays measuring the activity of the Zebrafish *Wnt3* promoter in HEK293T cells, co-transfected with hu∆β-cat, ZfSp5a, ZfSp5a-ΔDBD (left) or ZfSp5l1 (right). (F) Phylogenetic footprinting plot comparing 4kb genomic region encompassing the zebrafish *Wnt3* promoter with the corresponding regions of three teleost fish species. Pink peaks represent evolutionarily conserved modules as above; blue rectangles indicate regions of the *ZfWnt3* promoter tested for ZfSp5 binding in ChIP-qPCR assays. (G) ChIP-qPCR assays performed with cells co-transfected with ZfWnt3:Luc and the ZfSp5a expression construct. Note the significant enrichment in ZfSp5 but not ZfSp5ΔDBD in the PP1 and PP3 regions. Each data point in A, D and E represents one biological replicate. Statistical p-values: ∗≤ 0.05; ∗∗≤ 0.01; ∗∗∗≤ 0.001; ∗∗∗∗≤ 0.0001.

To further identify the sequences of the Wnt3-386 repressor involved in Sp5 repression, we tested four deletion constructs in HEK293T cells. We noticed that in all constructs *Wnt3* promoter activity is β-catenin inducible, while deleting different portions of the *HyWnt3* repressor results in an increase of the basal activity of the *HyWnt3* reporter construct. This suggests that different sequences of this region are required for an Sp5 independent repression of the *HyWnt3* promoter. Yet, the loss of repression observed in all these constructs was significantly lower to that observed when the whole repressive module was deleted, indicating that the different regions tested in this experiment act cooperatively to repress *Wnt3* expression. By contrast, Sp5 repression seems to require the two border regions (−386/−286 and −95/−1) of the Wnt3 repressive element since a significant reduction on luciferase activity upon co-transfection of β-catenin and Sp5 was only observed in the *HyWnt3-∆Rep-D3* construct (**Figure 5C, 5D**). This suggests that the *HyWnt3* repression by Sp5 requires the synergistic cooperation of two non-adjacent regions of the *Wnt3* promoter.

### Zebrafish Sp5 represses the activity of the zebrafish *Wnt3* promoter

To test whether the repressive activity of Sp5 also applies in vertebrates, we tested the activity of the zebrafish Sp5 transcription factor onto the zebrafish *Wnt3* promoter (3997 bp) in reporter assays and ChIP-qPCR experiments. As the HyWnt3-2149:Luc construct, the ZfWnt3-3997:Luc reporter shows a low basal activity, enhanced three fold by co-expressing hu∆β-cat (**Figure 5E**). The zebrafish Sp5 proteins, Sp5a and Sp5l1, reduce the activity of ZfWnt3-3997:Luc, a repressive effect that requires the Sp5 DNA-binding domain (**Figure 5E).** Although the Wnt3 promoter sequences of *Hydra* and zebrafish do not share obvious conserved stretches, a phylogenetic footprinting analysis comparing four teleost fish species revealed the presence of three evolutionary conserved modules in these *Wnt3* promoters (**Figure 5F**). ChIP-qPCR experiments using three pairs of primers along the *ZfWnt3* promoter identified the binding of ZfSp5a, but not ZfSp5aΔDBD, in two evolutionarily conserved modules located at the 5’ and 3’ ends (**Figure 5G**). Therefore, in zebrafish as in *Hydra,* Sp5 exerts a repressive effect on the *Wnt3* promoter through its DNA-binding activity.

**Figure 6.**
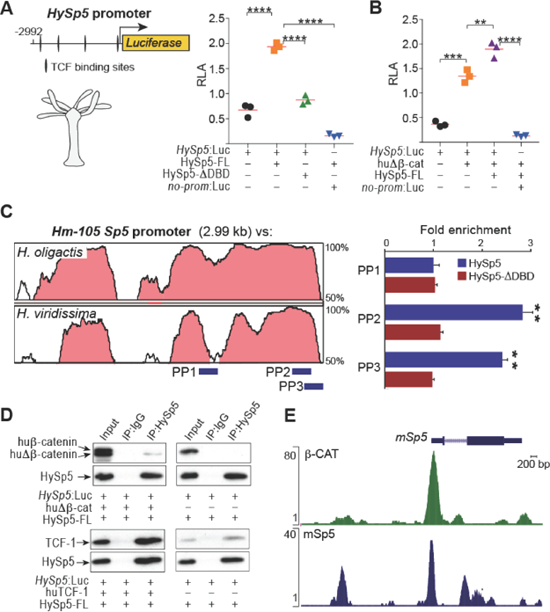
HySp5 activates its own promoter and interacts with TCF and β-catenin. **(A, B)** *HySp5* promoter activity measured in HEK293T cells that co-express HySp5 or HySp5-ΔDBD (A), or HySp5 together with huΔβ3-cat (B). Each data point in panels (A) and (B) represents one biological replicate. Statistical p-values were deduced from a student’s t-test: ∗∗ ≤ 0.01; ∗∗∗ ≤ 0.001; ∗∗∗∗ ≤ 0.0001. (C) Left: Phylogenetic footprinting plot comparing 2.99 kb genomic region encompassing the *Sp5* promoter in *Hm-105, H. oligactis* and *H. viridissima.* Pink peaks as in Figure 5. Blue rectangles: regions tested for Sp5 binding in ChIP-qPCR assays (right) performed in cells expressing HySp5:Luc in the presence of HySp5 or HySp5-ΔDBD. Note the significant enrichment in the PP2 and PP3 regions. (D) Immunoprecipitation (IP) of HySp5 co-expressed in HEK293T cells with huΔβ-catenin or huTCF-1. IP was performed with an anti-HA antibody and co-IP products detected with an anti β-catenln, or antl-TCF-1 antibody. Co-IPs were performed in two biological replicates. (E) ChIP seq profile showing the genomic occupancies of the mouse Sp5 and β-catenin over the genomic region encompassing the *Sp5* locus in mouse ES cells. The profiles were obtained by re-mapping publicly available dataset (Zhang et al. 2013; Kennedy et al. 2016). Note the overlap in the occupancies of Sp5 and β-catenin in the *Sp5* TSS vicinity.

### *Hydra* Sp5 acts through an autoregulatory loop

Since Sp5 can activate its own expression in mouse stem cells (Kennedy et al., 2016), we decided to test whether HySp5 also positively modulates the activity of its own promoter. We expressed HySp5-FLin HEK293T cells and measured a three fold increase in *HySp5-* 2992:Luc activity, an effect not observed when HySp5-ΔDBD is co-expressed **(Figure 6A)** but enhanced in the presence of hu∆β-cat (**Figure 6B**). This indicates that HySp5 enhances its own expression through promoter DNA-binding, possibly interacting with the β-catenin/TCF complexes that bind the TCF-binding sites present in the *HySp5* promoter. A phylogenetic footprinting analysis in three evolutionarily-distant *Hydra* species detected three conserved regions over the 3 kb of *Sp5* upstream sequences (**Figure 6C**, left panel). Next we designed primers mapping two of these regions to perform ChIP-qPCR experiments in HEK293T cells and observed a HySp5 enrichment in the region located immediately upstream of the *HySp5* predicted Transcriptional Start Site (TSS) (**Figure 6C**).

To elucidate whether HySp5 interacts with the TCFβ-catenin complex, we used HEK293T cells co-expressing HySp5 and hu∆β-catenin or HySp5 and huTCF-1 in co-immunoprecipitation assays. We detected interactions between HySp5 and β-catenin, as well as between HySp5 and TCF-1 (**Figure 6D** – **Figure Supplement 8A-B**), indicating that HySp5 can directly bind the TCFβ-catenin complex. We also noted that HySp5 interacts with the endogenous β-catenin and TCF-1 proteins. We then compared the occupancies of the mouse Sp5 and β-catenin proteins on the mouse *Sp5* promoter in ES cells (Zhang et al., 2013; Kennedy et al., 2016) and noted an overlap between their respective binding domains (**Figure 6E**), supporting the hypothesis that Sp5 auto-regulates its activity via β-catenin.

### Identification of DNA motifs bound by the *Hydra* and zebrafish Sp5 transcription factors

To identify the DNA motifs bound by *Hydra* and zebrafish Sp5, we performed ChIP-seq analyses in HEK293T cells expressing either HySp5 or ZfSp5a, and compared the results with a dataset reporting the genomic occupancies of β-catenin and Sp5 in mammalian cells (Zhang et al., 2013; Kennedy et al., 2016) (**Figure 7A**). Genome-wide, we identified 115’566 regions significantly enriched for HySp5, 331’89 for ZfSp5a, 106’220 of these being enriched for both, suggesting a large conservation of DNA-binding properties between *Hydra* and zebrafish Sp5 (**Figure 7B**, upper panel). However a high number of regions (224’869) is recognized by ZfSp5a only, indicating that vertebrate Sp5 transcription factors likely bind a spectrum of regulatory sequences broader than that of *Hydra* Sp5. Despite this difference, the gene assignment analysis revealed a similar number of genes potentially regulated by the HySp5 and ZfSp5a proteins, with 15’884 genes bound by both proteins (**Figure 7B**, bottom panel). Furthermore, we compared the spatial distribution of the regions bound by HySp5 and ZfSp5a relative to the TSS of their assigned genes and we found the HySp5 protein proportionally more frequently enriched in the close vicinity of the TSS (< 1 kb) than ZfSp5a that predominantly binds to mid-long distances, i.e. regions located between 50 and 100 kb from the TSS (**Figure 7C**).

The identification of motif consensus among the Sp5-enriched regions revealed the same core consensus motif GG(A/T/C)GG, centrally enriched within both the HySp5 and ZfSp5a bound sequences, very similar to the motif described for the human Sp5 protein (Huggins et al., 2017). Two other consensus motifs were found in the HySp5 dataset but neither in the ZfSp5a dataset, nor in the human or mouse Sp5 ChIPseq analyses (**Figure 7D – Figure Supplement 9**). Among these, the GGG(T/C)GTG motif is very similar to the motif recognized by the other Sp proteins (SP3: E=4.97e-03, q= 4.93e-03; SP8: E=2,8E-2, q=7,26e-3; SP1: E=2.93e-2, q=7,26e-3 SP4: E:6.43e-02, q= 1.28e-02) or by the KFL family members (e.g: Kfl1 : E=1.36e-03, q= 2.70e-03). This result suggests that HySp5 recognizes a more general Sp/Klf consensus binding site than ZfSp5a, while the vertebrate Sp5 proteins evolved towards a more restricted target sequence (GG(A/T/C)GG), although bound with a higher affinity. We used these predicted HySp5 and ZfSp5a consensus matrixes to identify putative Sp5-binding sites in the upstream sequences of *Wnt3* and *Sp5* in *Hydra* and zebrafish. As shown in **Figure Supplement 9,** evolutionarily-conserved although distinct putative Sp5-binding sites can be predicted in each context, largely overlapping with the regions enriched in the ChIP-qPCR assays, further supporting the idea that Sp5 directly binds and regulates its own promoter as well as that of *Wnt3.* In the case of the *HyWnt3* promoter, a putative Sp5 binding site was scored in the upstream region of the Wnt3 repressor, in agreement with the observation that the (−386 to −285) region is required for Sp5-dependent repression. Instead, no binding site was scored by this approach in the 100bp region immediately upstream of the *HyWnt3* Transcriptional Start Site (TSS). Another putative Sp5 binding site was scored in the central region, although this region was not sufficient per se for the Sp5 mediated *HyWnt3* promoter repression.

### Binding of *Hydra* and Zebrafish Sp5 onto the mouse and human genomic sequences

Next we focused on the human *SP5* and *WNT3* loci and their occupancies by HySp5 and ZfSp5 in HEK293T cells. The ChIP-seq data show a strong enrichment in HySp5 and ZfSp5a binding within the human SP5 promoter (**Figure 7E**), in agreement with the occupancies previously reported in mouse and human ES cells (Kennedy et al., 2016; Huggins et al., 2017). The binding profiles of HySp5 and ZfSp5a on the intronic and upstream regions of the human SP5 gene as well as in the upstream LOC440925 loci are highly related, further supporting the hypothesis of a functional equivalence between the *Hydra* Sp5 and the vertebrate cognate transcription factors. In tetrapods, the *Wnt3* gene is physically located in the vicinity of *Wnt9b,* while *Wnt3a* and *Wnt9a* are also clustered on the same chromosome (Garriock et al., 2007). The analysis of the binding of HySp5 and ZfSp5a in the human WNT3/WNT9b genomic region shows a strong enrichment for both proteins within the introns of both genes, as well as within the *WNT3/WNT9b* intergenic region, close to the *WNT9b* TSS. This result suggests that beside *WNT3,* Sp5 can regulate the expression of other *WNT* paralogs, at least in mammals (**Figure 7F**).

We also observed in human HEK293T cells an enrichment of HySp5 and ZfSp5a binding within the genomic region of several Wnt downstream target genes such as *Axin2, T-Bra, Lgr5,* in agreement with the binding of Sp5 in mouse ES cells (**Figure Supplement 10**). We also found *Hydra* Sp5 binding near the TSS of the *Polo-like Kinase 4 (PLK4)* gene, supporting the previous report showing that *PLK4* is a Sp5 target in human ES cells (Huggins et al., 2017). By contrast, we did not detect any Sp5 binding on the human *NANOG* sequences, a gene absent from cnidarian genomes (Watanabe et al., 2009). HySp5 seems to be able to bind a large number of Wnt/β-catenin downstream targets, even though the binding of Sp5 in mouse ES cells seems to differ, likely reflecting cell-type and/or species-specific differences in Sp5 occupancies. This role of Sp5 in the regulation of Wnt/β-catenin signaling is also supported by GO term enrichment analysis of putative Sp5 transcriptional targets, which identifies the WNT signaling pathway as among the three most enriched categories (**Figure 7G**).

**Figure 7.**
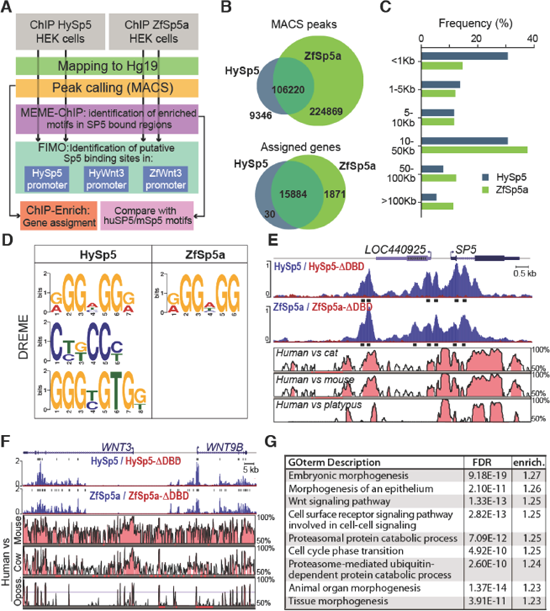
Genome-wide characterization of *Hydra* and zebrafish Sp5 binding sites in HEK293T cells. **(A)** Schematic workflow of the *in silico* analysis of the dataset from ChIP-seq experiments performed in HEK293T cells expressing HySp5 or ZfSp5a (see details in Materials and Methods). (B) Venn diagrams comparing the human genomic regions bound by HySp5 and ZfSp5a in the ChIP-seq experiments normalized to their respective input chromatin coverages (top panel) and the genes associated to the respective Sp5-enriched fragments (bottom panel). ZfSp5a binds approximately 3x more regions than HySp5 in the human genome, however the set of genes regulated by the two orthologs is similar. (C) Bar plot showing the frequency distribution of the genomic distances between the HySp5/ZfSp5a enriched regions and the transcription start site (TSS) of the genes to which they are assigned. (D) Weight matrixes of the sequences recognized by HySp5 and ZfSp5a respectively, deduced from the human genome-wide analysis of the regions found enriched in ChIP-seq experiments with the DREM motif discovery tool. (E, F) ChIP-seq experiment showing the genomic occupancies of the HySp5 and ZfSp5a proteins (blue) in the genome of human HEK293T cells expressing these proteins. Regions significantly enriched versus the control input chromatin (black rectangles below each track) were detected with the MACS2 tool. No enrichment is scored when HEK293T cells express Sp5 proteins lacking the DBD. Strong enrichment was detected in the upstream and intronic sequences of *SP5* (E), in the intronic sequences of *WNT3,* and in the promoter and intronic sequences of the neighboring *WNT9B* locus (F). Most Sp5-enriched sequences correspond to evolutionarily-conserved regions across mammals as shown by the phylogenetic footprinting analysis comparing the *SP5* (E), *WNT3* and *WNT9B* (F) genomic regions in four mammalian species (pink peaks). (G) Table summarizing the 10 most enriched GO terms associated with the genes assigned to the Sp5-enriched regions. GO term search was performed using the Gorilla software to compare the genes assigned to Sp5 bound regions in both HySp5 and ZfSp5a ChIP-seq experiments versus the full list of human genes.

## DISCUSSION

### *Sp5,* an evolutionarily-conserved target gene of Wnt/β-catenin signaling

Studies performed in developing vertebrates show that *Sp5* is a target of Wnt/β-catenin signaling as recorded in zebrafish (Thorpe et al., 2005; Weidinger et al., 2005), mice (Fujimura et al., 2007), *Xenopus* (Park et al., 2013), as well as in self-renewing mouse ESC (Kennedy et al., 2016) or differentiating human ESCs and iPSCs (Ye et al., 2016; Huggins et al., 2017). In line with these results, we show that in *Hydra, HySp5* is positively regulated by Wnt/β-catenin signaling as evidenced by its up-regulation when Wnt/β-catenin signaling is pharmacologically enhanced. These results illustrate the deep conservation of the Wnt/β-catenin dependent regulation of *Sp5* across eumetazoans. Wnt5, another candidate identified in the screen might also play a role in head inhibition, as a putative inhibitor of the canonical Wnt pathway (Mikels and Nusse, 2006; Nemeth et al., 2007) and a possible HySp5 target gene. By contrast, secreted Wnt antagonists such as dickkopf (Dkk) (Glinka et al., 1998) or Notum (Kakugawa et al., 2015), both expressed in *Hydra,* were not identified in this screen. Silencing *Notum* does not lead to a multiheaded phenotype (M.C.V. unpublished).

This study identifies the transcription factor Sp5 as a key inhibitory component of the *Hydra* head organizer as Sp5 fulfills the five criteria we initially fixed, derived from the predicted properties of the head inhibitor and from the previous identification of Wntβ-catenin signaling as the head activator (Nakamura et al., 2011). Sp5 globally fits the Turing Gierer-Meinhardt model as *HySp5* expression is controlled by Wnt3β-catenin signaling, graded along the body axis, reactivated during head regeneration, while HySp5 acts as a *Wnt3* repressor and represses ectopic head formation (**Figure 8**). However several features diverge from the expected properties for the head inhibitor.

### *Sp5* expression is globally consistent with a graded Sp5 activity along the body axis

In *Hydra, HySp5* exhibits an expression that is predominantly epithelial and graded from apical to basal along the animal axis (criteria-2), suggesting three levels of Sp5 activity in homeostatic conditions, maximal in the head, intermediate in the gastric region and null in the lower fourth. However this analysis of *HySp5* expression in intact animals also shows a lack of expression at the very apical tip of the hypostome, the region where *HyWnt3* expression, and most likely Wnt3 activity, is maximal. Two distinct cis-regulatory elements in the *Wnt3* promoter were previously identified, an activator and a repressor element, the latter restricting *HyWnt3* expression to the distal tip of the head (Nakamura et al., 2011). The *HySp5* pattern is thus consistent with the prediction that the inhibitor should be absent or should not bind the repressor element in this area. As *HySp5* is a direct target of Wnt3β-catenin signaling (criteria-1), an additional negative regulation has to take place in this most apical area, to prevent *HySp5* expression. This local regulation remains to be identified.

### Temporal constraints on *Wnt3* and *Sp5* up-regulations in head-regenerating tips are consistent with a rapid head organizer formation after bisection

*HySp5* is re-expressed early during head regeneration, although as expected, later than *Wnt3.* This temporal parameter is indeed essential for the establishment of a *de novo* head organizer as evidenced by transplantation experiments that accurately measured the successive re-activation of the two head organizer components, with head activation restored within 12 hpa while head inhibition comes back later, detectable at 24 hpa (MacWilliams, 1983a, 1983b). This differential timing is a necessary condition for a *de novo* head organizer formation. Here, we used the RNA-seq data to compare the respective regulations of *Wnt3* and *Sp5* in regenerating tips after decapitation or mid-gastric bisection. While *Wnt3* is rapidly up-regulated to reach a plateau value at 4 hpa, *HySp5* shows an initial drop in expression within the first two hours following bisection, then an up-regulation and a peak of expression detected at 8 hpa, four hours after that measured for *HyWnt3.* If one assumes that the reestablishment of active Wnt3 and Sp5 proteins follows similar kinetics, then this four hours time window corresponds to a period when Wnt3/β-catenin signaling is active but Sp5 still inactive as *Wnt3* repressor, leaving sufficient time to instruct tissues to form a head.

**Figure 8.**
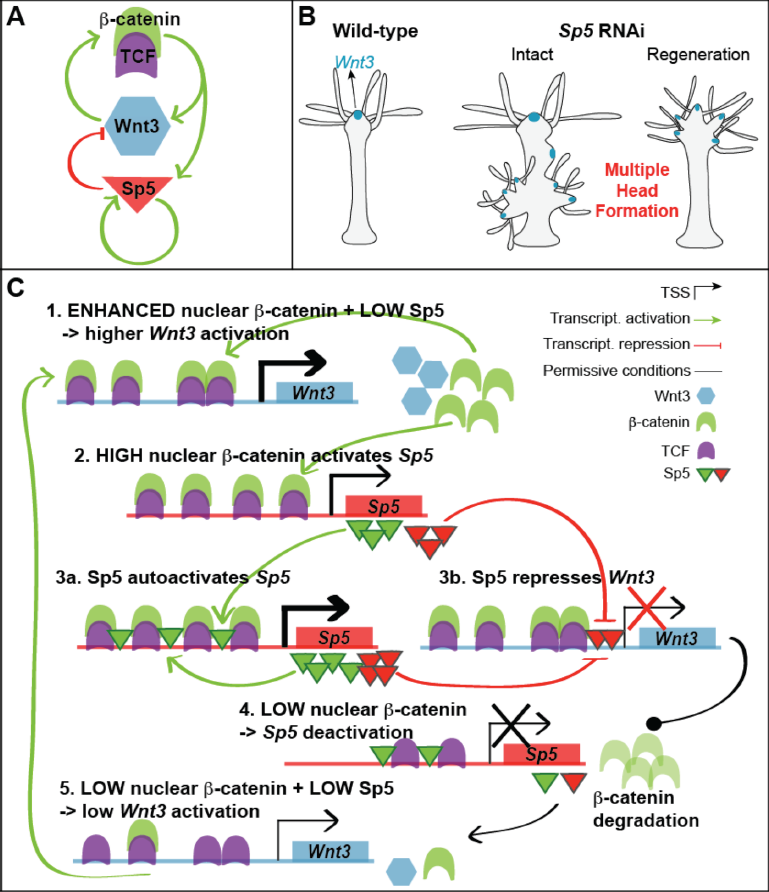
Working model of the feedback loop involving Wnt3/β-catenin-TCF and Sp5 transcriptional activities. **(A)** Schematic view of the feedback loop involving the growth factor Wnt3, the co-activator β-catenin, and the transcription factors TCF and Sp5. (B) Knocking down *HySp5* in intact and regenerating *Hydra* triggers multiple head formation. (C) Auto- and cross-regulatory interactions involving Wnt/β-catenin signaling that positively regulates *Wnt3* expression via β-catenin stabilization (steps 1 and 5, autoactivation) as well as *HySp5* expression (step 2). Once expressed, HySp5 positively auto-regulates its own expression, likely by interacting with the TCFβ-catenin complex (step 3a), but also represses *Wnt3* expression through the *Wnt3* repressor element (step 3b). As a consequence of *Wnt3* down-regulation, Wnt3 signaling and nuclear β-catenin are reduced, leading to a reduced *Wnt3* and *Sp5* expression (step 4). In the absence of Sp5, a low amount of Wnt3 suffices to auto-activate the *Wnt3* promoter (step 5). This feedback loop relies on the dual transcriptional activity of Sp5, positive on its own promoter (green triangles), repressive on the *Wnt3* promoter (red triangles), a property that appears conserved across eumetazoan evolution. A dual Sp5 regulation might apply to other *Wnt* genes and/or *Wnt* target genes. For temporal aspects of these regulations, see the discussion section.

### The transcription factor Sp5 acts as a novel repressor of Wnt3/β-catenin signaling

Sp5 from *Hydra* or from zebrafish inhibit Wnt/β-catenin signaling by repressing the *Wnt3* promoter activity. Both the reporter assays and the ChIP-qPCR experiments reveal that HySp5 directly binds the repressor element of the *HyWnt3* promoter. Our findings that zebrafish Sp5a and Sp5l1 also repress the activity of the zebrafish *Wnt3* promoter and that both *Hydra* and zebrafish Sp5 can bind the human Wnt3 promoter, further suggest that this mode of Wnt inhibition originated early in metazoan evolution and was maintained across eumetazoans. In fact, in mammals, Sp5 was identified as a repressor of Sp1 target genes in the mouse brain (Fujimura et al., 2007) and as a terminator of the transcriptional programs initiated by Wnt in human pluripotent stem cells (Huggins et al., 2017). However it is as yet unclear whether human SP5 directly represses *Wnt3* promoter activity. A recent study shows as a side observation that the level of *Wnt3* expression is increased in human ESCs knocked-out for SP5 (Huggins et al., 2017), supporting this hypothesis. Furthermore, our ChIP-seq data demonstrate that both *Hydra* and zebrafish Sp5 bind the human WNT9B promoter, suggesting that the repressor effect of SP5 proteins is not restricted to *WNT3.* Although further studies are required to demonstrate the *Wnt3* repressing effect of Sp5 *in vivo,* this study illustrate that the antagonism between Sp5 and Wnt3 is a highly conserved feature across eumetazoans.

### Sp5 might act autonomously and/or regulate the production of diffusible inhibitors

These results support a scenario where HyWnt3 acts as a short-range activator to sustain its own activity in the head organizer, while Sp5 would prevent *Wnt3* expression over long distances. The Gierer-Meinhardt model predicts that the head inhibitor is a diffusible substance acting non-cell autonomously across the tissue layers. As a transcription factor, HySp5 is preferentially not diffusible, acting cell-autonomously. However transcription factors can be secreted, as reported for the helix-turn-helix transcription factor EspR in bacteria (Raghavan et al., 2008) or for some homeoproteins that exert non-cell autonomous functions in the mammalian brain (Bernard et al., 2016). Also, Sp5 might up-regulate target genes that encode secreted peptides or proteins with the expected head inhibitory functions. Such target genes, possibly taxon-specific, remain to be identified.

Finally, in *Hydra*, the presence of diffuse inhibitors might not be necessary for head inhibition, as long as *Wnt* and *Sp5* genes are co-expressed, which is the case at least in the body column. Also, Wnt signals, which are numerous to be emitted from the apical region, might also act over long-range distances to activate *HySp5* expression with lipid-binding proteins or cytonemes modulating the spread of Wnt proteins as observed in *Drosophila, Xenopus* and zebrafish (Mii and Taira, 2009; Mulligan et al., 2012; Stanganello et al., 2015). The inhibition of Wnt3 signaling along the *Hydra* body axis might thus be solely mediated by transcriptional repression, with Sp5 auto-regulating its own expression and tightly tuning the level of Wnt signals, with or without the contribution of Wnt/βcatenin signaling.

### The *Sp5* RNAi phenotype highlights the head inhibitory function of HySp5 and its dynamic regulation

Consistent with its *Wnt3* repressor function, *HySp5* silencing triggers the formation of multiple heads along the body column of both intact and head-regenerating animals (**Figure 8**). This phenotype is different from the ones obtained with pharmacological treatments, either with the GSK3-β inhibitor ALP (Broun et al., 2005; Augustin et al., 2006; Guder et al., 2006) or the recombinant Wnt3 that directly promotes β-catenin signaling (Chera et al., 2009; Lengfeld et al., 2009), where ectopic tentacles form first, and heads appear several days later. Here the knock-down of *HySp5* leads to the direct and rapid formation of fully functional ectopic heads, which preferentially form in the budding zone, a developmental competent region in adult animals where the expression of both *Wnt3* and *β-catenin* is quite dynamically regulated (Hobmayer et al., 2000; Lengfeld et al., 2009). By increasing the number of electroporations, we also noted the formation of ectopic heads in the apical half. However the development of these heads was incomplete. Similarly we never observed supernumerary heads at the apex of homeostatic HySp5(RNAi) animals. These results likely reflect a too partial and/or too transient silencing in regions where *HySp5* expression is twice higher than along the mid-gastric region. Therefore, we interpret the homeostatic HySp5(RNAi) phenotype as the consequence of the short time window(s) when HySp5 activity is lowered in homeostatic tissues that have the highest potential for setting up an organizer. Indeed *HySp5* expression is seen as highly dynamic and a drop of HySp5 protein level rapidly induces a derepression of *Wnt3* expression, which leads to an up-regulation of β-catenin activity, and in turn to the up-regulation of *Wnt3* expression followed by that of *Sp5* (**Figure 8**). The oscillatory nature of *Sp5* expression in the head organizer remains to be explored but is fully compatible with an auto-regulatory loop involving two transcription factors (Widder et al. 2009).

### HySp5 autoregulates its expression, likely through direct interactions with TCF and/or β-catenin

As another divergence with the Gierer-Meinhardt model, we found that HySp5 activates its own promoter. Both the reporter assays and the ChIP-qPCR data demonstrate that HySp5 directly binds its own promoter, while the ChIP-seq data also suggest that HySp5 is able to bind the human SP5 promoter. These observations are consistent with studies showing that the mouse and human Sp5 proteins directly bind and activate their own promoter (Kennedy et al., 2016; Huggins et al., 2017). In addition, HySp5 does enhance the activating effect of β-catenin on the *HySp5* promoter, likely through direct interaction with TCF-1 and/or β-catenin as observed *in vitro.* A recent study demonstrates a direct interaction between the zinc finger domain of mouse Sp5 and the HMG domain of TCF-1 and LEF-1, while no direct interaction was observed for β-catenin (Kennedy et al., 2016). Also the formation of active TCF/LEF-β-catenin complexes appears necessary for Sp5 DNA-binding in mouse ESCs (Kennedy et al., 2016). The interactions between HySp5 and TCF/β-catenin remain to be explored *in vivo.*

In summary, we favor a model where Wnt/βcatenin signaling up-regulates *Sp5* expression, a step followed by the binding of Sp5 onto its own promoter together with the TCF/β-catenin complex to robustly activate its own expression, while in parallel Sp5 binds to the *Wnt3* promoter to repress its expression (**Figure 8**). Therefore in *Hydra* HySp5 acts in a feedforward loop as recently suggested for mouse Sp5 (Kennedy et al., 2016). Besides elucidating the molecular nature of the head inhibitory component in the hydrozoan *Hydra,* we identified Sp5 as a novel, evolutionarily-conserved repressor of *Wnt3* gene expression. This finding has a potential relevance for cancer biology, as beside its repressor function in developmental and regenerative contexts, Sp5 might also be used to play a negative role in oncogenic contexts where Wnt/βcatenin signaling is enhanced.

## MATERIALS AND METHODS

### Animal culture and drug treatment

All experiments were carried out with *Hydra vulgaris (Hv)* from the Basel, AEP or *Hm-105* strains. Cultures were maintained in Hydra Medium (HM: 1 mM NaCl, 1 mM CaCl2, 0.1 mM KCl, 0.1 mM MgSO4, 1 mM Tris pH 7.6) or in Volvic water, supplemented with 0.5 mM CaCl_2_. Animals were fed two to three times per week with freshly hatched *Artemia* nauplii and starved for four days before any experiment. For drug treatments Hv_Basel were treated for two days with 5 μM Alsterpaullone (Sigma) diluted in HM, 0.015% DMSO (Sigma), then rinsed 3x in fresh HM and used for RNA extraction or *in situ* hybridization experiments.

### RNA-seq data

The procedure for producing RNA-seq data is described in (Buzgariu et al., 2018). A detailed manuscript about the analysis of the transcriptomes used in this study is in preparation. It will provide a full release of the data and codes (Wenger et al. in preparation).

### Multiple sequence alignment and phylogenetic analyses

The multiple sequence alignment was generated using T-Coffee (Notredame et al., 2000). The conserved zinc finger domains of Sp5 were identified using PROSITE (Sigrist et al., 2002) and the SP and Btd boxes were identified as described in (Bouwman and Philipsen, 2002) and visualized by IBS (Liu et al., 2015). For the phylogenetic analysis of the Sp5, Sp-related and Klf-related gene families, sequences from *Hydra* as well as from species representative for cnidarian, ecdysozoans, lophotrochozoans and deuterosomes were retrieved from Uniprot or NCBI, aligned with Muscle align (www.ebi.ac.uk/Tools/msa/muscle/) (Edgar, 2004a, 2004b) and tested in iterative PhyML 3.0 analyses using the LG substitution model, 8 substitution rate categories and 100 bootstraps (Guindon et al., 2010).

### Plasmid constructions

To generate the *HyWnt3:Luc* construct 2’149 bp of the *Hydra Wnt3* promoter were transferred from the hoTG-HyWnt3FL-EGFP construct (kind gift from Thomas Holstein, Heidelberg) (Nakamura et al., 2011) into the pGL3 reporter construct, a kind gift from Zbynek Kozmik, Prague (Fujimura et al., 2007). For the HyWnt3ΔRep:Luc construct, the whole HyWnt3:Luc plasmid sequence was PCR-amplified except the 386 bp corresponding to the repressor element. For the *ZfWnt3*:Luc construct 3’997 bp of the zebrafish *Wnt3* promoter were transferred from pEGFP-Wnt3 (kind gift of Cathleen Teh, Singapore) (Teh et al., 2015) into pGL3. For the HySp5:Luc construct, 2’992 bp of the *Hydra Sp5* promoter were PCR-amplified from *Hm-105* genomic DNA and subcloned into pGL3. To express HA-tagged HySp5, ZfSp5a, ZfSp5l1 proteins, a C-terminal HA-tag was introduced into the pCS2+ constructs encoding the *Hydra Sp5* (human codon-optimized), zebrafish *Sp5a* and *Sp5l1* full-length coding sequences. For the constructs expressing truncated proteins (Figure 3C): HySp5-ΔSP construct was produced by inserting a human codon-optimized *HySp5* sequence lacking 110 amino acids of the N-terminal end together with a C-terminal HA-tag into pCS2+. The HySp5-βDBD and HySp5-βSP-βDBD constructs were generated using the QuikChange Lightning Multi Site-Directed Mutagenesis Kit (Agilent Technologies) following the manufacturer’s instructions. To generate the ZfSp5a-βDBD construct, the entire ZfSp5a plasmid sequence except the region encoding the DNA-binding domain was PCR-amplified and self-ligated. For in situ detection of transcripts, the *HyWnt3, HyBra1* and *HySp5* PCR products were cloned into pGEM-T-Easy (Promega). All constructs were verified by sequencing. All plasmids are listed in **Table Supplement 4** and primer sequences in **Table Supplement 3A**.

### RNA interference

RNA interference (RNAi) was induced by electroporating a mix of Sp5 or control siRNAs (Sp5A/B/C, scrambled see **Table Supplement 3B**) using the Biorad GenePulser Xcell electroporation system as reported by (Watanabe et al., 2014). In short, intact *Hydra (Hv_Basel)* were briefly washed and incubated for 45 min in Milli-Q water, treated for 5 minutes with 1.5% Bisolvon (Boehringer Ingelheim) and washed again briefly in water. 20 animals per condition were placed in 200 μl 10 mM sterilized HEPES (pH 7.0), then transferred into a 0.4 cm gap electroporation cuvette (Cell Projects Ltd) and 4 μM siRNA added. The cuvette was tapped 5-10 times to distribute the animals before inserting it into the shocking chamber The electroporation was carried out after 5 minutes under the following conditions: Voltage: 150 Volts; Pulse length: 50 milliseconds; Number of pulses: 2; Pulse intervals: 0.1 second. Immediately after electroporation, 200 μl of restoration medium (RM: 1.2 mM CaCl2, 0.24 mM MgSO_4_, 0.72 mM KCl, 2.5 mM TES, 1.2 mM Na pyruvate, 1.2 mM Na citrate, 1.2 mM glucose, 10 mg/l Rifampicin, pH 6.9) was added into the cuvette. The animals were then transferred into a 6-well plate containing 5 ml RM per well, which was replaced by HM 24 hours later.

### qPCR

Total RNA was extracted using the E.Z.N.A.^®^ Total RNA kit (Omega) and cDNA synthesized using the qScript™ cDNA SuperMix (Quanta Biosciences). qPCR was performed in a 96-well format using the SYBR™ Select Master Mix for CFX (Thermo Fisher Scientific) and a Biorad CFX96™ Real-Time System. TATA-box binding protein (TBP) was used as an internal reference gene. The relative *HySp5* expression level was calculated as described in (Pfaffl, 2001). Primer sequences for HySp5 and TBP can be found in **Table Supplement 3C**. For ChIP-qPCR, DNA was prepared as described in the ChIP-seq section and qPCR was performed as described above. Detailed primer sequences for the *HySp5* and *ZfWnt3* promoters are shown in **Table Supplement 3D**.

### Whole mount *In Situ* Hybridization (WISH)

*Hydra* were relaxed in 2% urethane/HM for one minute, fixed in 4% PFA prepared in HM (pH 7.5) for 4 hours at RT and stored in MeOH at −20°C for at least one day. Samples were rehydrated through a series of EtOH/PBSTw (PBS, Tween 0.1%) washes (75%, 50%, 25%) for 5 minutes each, washed 3x with PBSTw for 5 minutes, and treated with 10 μg/mL Proteinase K/0.1% SDS/PBSTw (Roche) for 10 minutes. The Proteinase K digestion was stopped by adding Glycine/PBST (4 mg/mL) and incubation for 10 minutes. The animals were washed 2x with PBSTw for 5 minutes, treated with TEA (0.1M) for 2× 5 minutes, followed by adding two times acetic anhydride to the TEA solution and incubation for 5 minutes (final concentration 0.25% (v/v) and 0.5% (v/v), respectively). The samples were then washed 2x with PBSTw for 5 minutes, post-fixed in 4% formaldehyde/PBSTw for 20 minutes, washed 4× with PBSTw for 5 minutes before adding the pre-warmed prehybridization buffer (PreHyb: 50% Formamide, 0.1 % CHAPS, 1x Denhardt’s, 0.1 mg/mL Heparin, 0.1% Tween, 5× SSC) and incubated for 2 hours at 58°C. In-between, 350 μL hybridization buffer (PreHyb with 0.2 mg/mL t-RNA and 5% Dextran) with 200 ng of DIG-labeled probe was heated for 5 minutes at 80°C and then placed on ice for 2 minutes. This mix was added onto the samples, then incubated for 19 hours at 58°C. Next the samples were rinsed 3x in pre-warmed PostHyb 1 (50% formamide, 5× SSC) and incubated for 10 minutes at 58°C in PostHyb 1, PostHyb 2 (75% PostHyb1, 25% 2× SSC; 0.1% Tween), PostHyb 3 (50% PostHyb1,50% 2x SSC; 0.1% Tween) and PostHyb 4 (25% PostHyb1,75% 2× SSC; 0.1% Tween). Samples were then washed in 2× SSC, 0.1% Tween and 0.2x SSC, 0.1% Tween (2× 30 minutes each). Afterwards, the samples were washed in Buffer I (1x MAB; 0.1% Tween) for 2× 10 minutes before blocking in 20% sheep serum/ Buffer I (Buffer II) for 1 hour and incubation with an anti-DIG-AP antibody (1:4000, Roche) in Buffer II over night at 4°C. On the following day, the animals were washed in Buffer I for 4x 15 minutes, then in NTMT (NaCl 0.1 M, Tris-HCl pH 9.5 0.1 M, Tween 0.1%) for 5 minutes and finally in NTMT, levamisole 1 mM for 2× 5 minutes. The colorimetric reaction was started by adding 500 μL staining solution (Tris-HCl pH 9.5 0.1 mM, NaCl 0.1 mM, PVA 7.8%, levamisole 1 mM) containing NBT/BCIP (Roche). The background color was removed by a series of washes in EtOH/PBSTw (30%/70%, 50%/50%, 70%/30%, 100% EtOH, 70%/30%, 50%/50%, 30%/70%), then in PBSTw 2× 10 minutes. Samples were post-fixed for 20 minutes in FA 3.7% diluted in PBSTw, then washed in PBSTw 3× 10 minutes and mounted with Mowiol. All steps were performed at RT, unless indicated otherwise.

### Whole mount immunohistochemistry

*Hydra* were fixed and stored in MeOH as described above, rehydrated through a series of MeOH/PBS washes (75%, 50%, 25%) for 15 minutes, washed 4x 10 minutes in PBS, then in PBSTr (PBS with 0.5% Triton-X100) for 45 minutes and incubation in 2% BSA/PBSTr for 1 hour. Samples were then incubated overnight in anti-RFamide antibody (Grimmelikhuijzen and Graff, 1985) (kind gift of C. Grimmelikhuijzen, 1:1000) at 4°C, washed 6x 10 minutes with PBSTr and incubated in anti-rabbit Alexa488 antibody (1:600, A21206, ThermoFisher Scientific) for 4 hours. Samples were then again washed 6x 10 minutes with PBSTr, incubated for 20 minutes in DAPI (1:5000, Roche), washed 2x 10 minutes with PBSTr and mounted with Mowiol. All steps were performed at RT, unless indicated otherwise.

### Generation of the AEP HyWnt3:GFP-HyAct:dsRED transgenic strain

To induce gametogenesis *H. vulgaris* of the strain AEP were fed 7x per week, then 1x per week, and 7x per week with freshly hatched *Artemia* nauplii. Thereafter, male and female animals were cultured together, resulting in fertilized embryos. The hoTG-HyWnt3FL-EGFP plasmid (kind gift from Thomas Holstein, Heidelberg) (Nakamura et al., 2011) was injected into one-cell stage embryos. Out of 504 injected eggs, 104 embryos hatched and 7/104 embryos exhibited GFP fluorescence in the hypostome.

### Mammalian cell culture

HEK293T cells, a kind gift from Ariel Ruiz i Altaba (Geneva Medical School), were maintained in DMEM High Glucose, 1 mM Na pyruvate, 6 mM L-glutamine, 10% fetal bovine serum. For the luciferase assays HEK293T cells were seeded into 96-well plates (5000 cells/well) and transfected 18 hours later with X-tremeGENE^TM^ HP DNA transfection reagent (Roche). To measure Firefly and *Renilla* luciferase activities, the samples were prepared using the Dual-Luciferase Reporter Assay System (Promega), transferred to a white OptiPlate^TM^-96 (PerkinElmer) and measured with a multilabel detection platform (CHAMELEON™). The plasmids listed in **Table Supplement 4** were transfected in HEK293T cells as follows:pGL4.74[hRluc/TK] (Promega): 1 ng, luciferase reporter constructs: 40 ng, CMV:huΔβ-cat: 10 ng Sp5 expression constructs: 20 ng, huWnt3 and huLRP6: 40 ng. Total DNA amount was adjusted with pTZ18R to 100 ng per well. The following plasmids were kindly given to us: pcDNA-Wnt3 (huWnt3) by M. Waterman (Addgene #35909) (Najdi et al., 2012), pcDNA6-hLRP6-v5 (huLRP6) by B. Williams (Holmen et al., 2002), pFLAG-CMV-human-β-cateninΔ45 (huΔβ-cat) by A. Ruiz i Altaba (Melotti et al., 2014).

### ChIP-seq

HEK293T cells were seeded into a 10 cm dish (920’00 cells) and transfected as described above with the HySp5-420, ZfSp5-377, HySp5ΔDBD-337, ZfSp5-ΔDBD-289 plasmids (3’666 ng). 24 hours later, cells were collected by scraping, washed twice in pre-warmed culture medium, fixed in 1% FA solution (Sigma) for 15 minutes. The fixative was quenched by adding Glycine (125 mM) and incubation for 3 minutes. The cells were washed once in ice-cold PBS and re-suspended in 5 mL chromatin prep buffer (Active Motif), containing 0.1 mM PMSF and 0.1% protease inhibitor cocktail (PIC). To release the nuclei the sample was transferred into a pre-cooled 15 mL glas Douncer and dounced with 30 strokes. After 10 minutes incubation on ice and centrifugation at 1250 g for 5 minutes at 4°C, the nuclei were re-suspended in 500 uL sonication buffer (1% SDS, 50 mM Tris-HCl pH 8.0, 10 mM EDTA pH 8.0, 1 mM PMSF, 1% PIC). Following another 10 minutes incubation on ice, the chromatin was sonicated with a Bioblock Scientific VibraCell 75042 sonicator (Amplitude: 25%, Time: 12 minutes, 30 seconds on, 30 seconds off, 24 cycles). Note: The sonication conditions were optimized to have a fragmentation size of around 250 bp. Then 100 uL of the sonicated chromatin was added to 900 uL ChIP dilution buffer (0.1% NP-40, 0.02 M HEPES pH 7.3, 1 mM EDTA pH 8.0, 0.15 M NaCl, 1 mM PMSF, 1% PIC) and incubated with 4 ug anti-HA antibody (NB600-363, Novus Biologicals) over night at 4°C on a rotator. Next the sample was loaded on a ChIP-IT ProteinG Agarose Column (Active Motif) and incubated for 3 hours at 4°C on a rotator. The column was washed 6 times with 1 mL buffer AM1 (Active Motif) and the DNA eluted with 180 uL of pre-warmed buffer AM4 (Active Motif). The sample was decrosslinked by adding 30 uL 10x TE buffer, 18 uL 5 M NaCl, 57 uL H2O and incubated for 5 hours at 65°C. 5 uL of RNAse A (10 ug/uL) was added and the sample incubated at 37°C for 30 minutes before adding 10 uL of Proteinase K (10 ug/uL), and further incubation for 2 hours at 55°C. The DNA was purified with the MiniElute PCR purification kit (Qiagen). For preparing the Input DNA, 5 uL sonicated chromatin was added to 5 uL 5M NaCL in 40 uL H2O, and incubated for 15 minutes at 95°C. Next the sample was incubated at 37°C for 5 minutes with 2.5 uL of RNAse A (10 ug/ul), 2.5 uL PK (10 ug/uL) was then added, and the incubation continued at 55°C for 30 minutes. 10 uL were taken for purification (MiniElute PCR purification kit from Qiagen).

### ChIP-seq data analysis

Demultiplexed ChIP-seq reads from our sequenced samples were mapped onto the Human GRCh37 (hg19) genome assembly using bowtie2, version 2.2.6.2 (Langmead and Salzberg, 2012). Significantly enriched regions were identified using MACS2 REF: (Zhang et al., 2008) (version 2.1.0.20151222.0). Coverage files were normalized by the millions of mapped reads in each sample using a manually created R script, and visualized with UCSC genome browser. Fasta formatted files containing 100 bp long sequences of significantly enriched regions from this analysis were obtained using the Table browser function from the UCSC browser. These files were used to identify enriched motifs for transcription factor binding sites using the MEME-ChIP Suite (Bailey et al., 2009) (http://meme-suite.org/tools/meme-chip) in discriminative mode. MACS2-enriched region form the Total Imput Chromatin controls of each experimental condition were used as reference. In both ChIP-seq experiments performed from HEK cells transfected with the HySp5 or ZfSp5 coding plasmids, significantly enriched motifs were identified and compared to previously described TF weight matrixes form the JASPAR CORE 2014 database (Bailey et al., 2009) using the TOMTOM tool of the MEME-ChIP suite. Significantly enriched motifs were used to scan the HySp5, HyWnt3 and ZfWnt3 promoters, using the FIMO tool (http://meme-suite.org/tools/fimo) (Grant et al., 2011) to identify putative Sp5 binding sites. Datasets of the SP5 (Huggins et al., 2017) and β-catenin (Estaras et al., 2015) genome wide occupancies in human ES cells were downloaded from GEO subseries GSM2756639 and GSM1579345 respectively. CHIP-seq datasets for the Sp5 and β-catenin occupancies in mouse ES cell (Kennedy et al., 2016) were downloaded from the GEO subseries GSE72989 and GSM1065517 respectively. All these datasets were re-mapped and analyzed following the same workflow used for our ChIP-seq experiments.

### Co-immunoprecipitation (Co-IP) assay and Western blotting

HEK293T cells were seeded into a 10 cm dish (920 000 cells) and transfected with *HySp5:Luc* (7320ng), hu∆β-cat (1830ng), huTCF-1 (1830ng) and HySp5 (3660ng). 24 hours later, Co-IP samples were prepared using the nuclear complex Co-IP kit from Active Motif. In short, cells were collected by scraping, centrifuged at 430x g for 5 minutes at 4°C, washed twice with 4 mL ice-cold PBS/Phosphatase inhibitors. The pellet was resuspended in 500 uL hypotonic buffer, incubated for 15 minutes on ice until 25 uL detergent was added, centrifuged at 14’000x g for 30 seconds at 4°C, resuspended in 100 uL complete digestion buffer, 0.5 uL enzymatic shearing cocktail added and incubated for 90 minutes at 4°C. Afterwards, 2 uL of 0.5 M EDTA was added, the sample incubated for 5 minutes on ice, centrifuged at 14’000x g for 10 minutes at 4°C and the supernatant transferred into a prechilled 1.5 mL tube. 100 ug extract was diluted to 500 uL with IP incubation buffer and 4 ug of an anti-HA antibody (NB600-363, Novus Biologicals) added and incubated over night at 4°C on a rotator. Note: 4 ug of rabbit IgG (12-370, Merck Millipore) was used as a control immunoglobulin. On the next day, the IP reaction was loaded on a Protein G Agarose column (Active Motif) and incubated for 1 hour at 4°C on a rotating wheel. The column was washed 3 times with 500 uL of ice-cold IP wash buffer supplemented with 1 mg/mL BSA and 3 times with 500 uL of ice-cold IP wash buffer supplemented with 300 mM NaCl. The column was centrifuged at 1250x g for 3 minutes at 4°C and 25 uL of 2x reducing buffer directly added onto the column. After incubation for 1 minute and centrifugation at 1250 g for 3 minutes at 4°C, 5 uL of pure glycerol (Sigma) was added and the sample boiled for 5 minutes at 95°C. The sample was then loaded on a 8% SDS-PAGE gel, electrophoresed and transfered onto a PVDF membrane (Bio-Rad). The membrane was blocked with 5% milk in TBS-Tw (TBS containing 0.1% Tween) for one hour until primary antibodies diluted in TBS-T 5% milk (1:2000) were added for ON incubation at 4°C: anti-HA antibody (NB600-363, Novus Biologicals), antiβ-catenin antibody (610153, BD Biosciences) and anti-TCF-1 (sc-271453, Santa Cruz Biotechnology). The next day, the membrane was washed 4x 10 minutes with TBS-Tw, incubated in anti-rabbit (ab99697, Abcam) or anti-mouse (W402B, Pomega) IgG horseraddish peroxidase antibody (1:5000) for one hour. The protein signals were visualized with the Western Lightning^®^ Plus-ECL reagent (PerkinElmer). Note: 10 ug of extract were used as the Input sample. All steps were performed at RT unless indicated otherwise. The human TCF1 plasmid was a gift from Kai Ge (Addgene plasmid # 40620) (Wang et al., 2012).

### Sp5 expression in zebrafish embryos

For all zebrafish experiments, colonies of the strain AB-Tu or Nacre were used, with animals maintained at 28°C with a maximal density of five fish per liter in a 14 hour light-10 hour dark cycle. The fish were fed twice a day with 2-day-old Artemia and fish embryos incubated at 28°C. For overexpression experiments, capped sense mRNAs were synthesized using the mMESSAGE mMACHINE^®^ Transcription Kit from Ambion (Ambion, Austin, TX USA) and 400 pg of *HySp5, HySp5ΔDBD, HySp5ΔSP* or *HySp5-ΔSPΔDBD* mRNAs injected into one cell stage embryos. For mRNA co-injection experiments, injected amounts were as follows: 400 pg of HySp5 and 4 pg of ZfWnt8 mRNA. All embryos were scored for phenotypes 48 hours post fertilization. ZE14 pCS2P+ wnt8 ORF1 was a gift from Randall Moon (Addgene plasmid # 17048) (Lekven et al., 2001).

### Statistical analysis

All statistical analyses were performed with the software GraphPad Prism7.

### *Hydra* genomes

Five clonal animals of the species *Hydra viridissima* and *Hydra oligactis* were sampled independently to extract DNA material using the DNeasy Blood & Tissue kit (Qiagen). Sequencing libraries were prepared using the TruSeq Nano DNA kit (Illumina), with 350bp insert sizes, and sequenced paired-end using 150 cycles on an Illumina HiSeq X Ten sequencer by Macrogen Inc. Average and standard deviations of insert sizes of the sequenced reads were measured using 10 mio reads mapped to a preliminary assembly of each genome, then the two genomes were assembled using MaSuRCA v3.2.1 (Zimin et al., 2013). All scaffolds, and unplaced contigs of more than 500 bp were retained in the final set of sequences. The redundancy of each assembly was reduced by using CD-HIT-est v4.7 (Li and Godzik, 2006) with a 100% identity threshold.

### Accession numbers

The *Hydra Sp5* sequence was deposited at GenBank (MG437301). The genome assemblies and reads are available under the BioProject PRJNA419866. ChIP-seq experiments have been deposited with the GEO database under the following accessions: GSE110277.

## ACKNOWLEDGEMENTS

This work was supported by the Canton of Geneva, the Swiss National Foundation (SNF 31003A_149630), the Human Frontier Science Program (grant no. RGP0016/2010), the NCCR ∘Frontiers in Genetics', the Claraz donation and the de Staël foundation. The authors thank Denis Duboule for valuable input on the manuscript, Carol Gauron for excellent technical assistance, Nenad Suknovic for help with image acquisition and the IGe3 Genomic Platform for ChIP-seq library preparation and sequencing.

## Author contributions

M.C.V. performed *Hydra* and cell culture experiments, performed biochemical assays and prepared ChIP-seq samples; L.B. analyzed ChIP-seq data. M.C.V. and L.B. performed ChIP-qPCRs. L.I.O. contributed to plasmid constructions and *in situ* hybridizations; C.R. and S.V. performed zebrafish experiments; Y.W. and B.G. designed the high-throughput transcriptomics, Y.W. produced and processed the high-throughput transcriptomics as well as the whole genome data; C.P. produced the transgenic line; M.C.V. and B.G. conceived the study, M.C.V., L.B. and B.G. wrote the manuscript; B.G. supervised the study and obtained the funding.

